# Biotransformation of agro-industrial waste to wealth in Africa through mushroom farming

**DOI:** 10.1101/2021.03.01.433330

**Authors:** Victor S. Ekun, Clementina O. Adenipekun, Olufunmilayo Idowu, Peter M. Etaware

## Abstract

Discarding Waste in open drainages, water bodies and unauthorized dumpsites is a common practice in Africa that is deleterious to the global ecosystem. Global warming, flooding and land encroachment are direct impact of these actions. Therefore, a proposed eco-friendly solution to efficient waste management in Africa is “Mushroom Farming”. The region between Latitude 6.43–7.95°N and Longitude 2.88–5.24°E (Nigeria) was the research focus. Mushroom samples were collected, identified, coded and geo-tagged in the field, while Laboratory analyses and mushroom cultivation was conducted in NIHORT and University of Ibadan. The result showed that postharvest waste (rice straw) was best for spawning (Ek1-8→23days, LA1-8→21days, OG1-8→20days, and OY1-8→22days), while agro-waste from *Gliricidium sepium* facilitated early pin head emergence (26days) and fruiting (28days). Also, agro-waste from *Cedrela odorata* facilitated early maturation (3days after fruiting), and those from *Mangifera indica, G. sepium* and *C. odorata*, improved yield (9.05g), pileus size (4.24g) and dry matter (3.01g), respectively. The Laboratory analyses showed that LA1-8 had the best N_2_, Na and Ca contents (13.59, 70.49 and 61.90mg/100g, respectively), while fat, fibre and carbohydrate contents were highest in samples from EK1-8 (6.60%), OS1-8 (25.13%) and OG1-8 (54.23%), respectively. Mushroom farming is the key to efficient transformation of waste to wealth in Africa.

## 1.0 Introduction

The constant deposition of waste in open drainages, water bodies and unauthorized dumpsites widely practiced in Africa is a course for concern [1]. Waste management is a theme that needs global redress, as this problem is not synonymous to Africa alone. Sadly, Africa is at the forefront of continents heavily populated by untreated waste. The impact of untreated waste in Africa is very severe, as it is partly responsible for the dysfunction of the ecosystem, unfavourable climate change, global warming and irregular fluctuation in weather patterns over the last century etc. [1]. Also, the rapid loss of productive land useful for agriculture and the increase in the incidence of flooding recorded over the past decade in Africa, are products of land encroachment by waste, blocked drainages, increased soil acidity and nutrient depletion caused by increased microbial activities in the soil [1].

Chemical analysis of waste showed that irrespective of their source of production (Domestic, Agro-Industrial and Industrial), they contain varying amount of organic compounds, inorganic and radioactive elements. Some of the components of waste are recalcitrant, insoluble in water and non-degradable e.g. polyethylene, lignocellulose, PAHs, PCBs, CFCs, radioactive elements, etc. [2] Some of these wastes can be used as resource for commercial production of food and value-added food products like mushroom cultivation [2], while others can be effectively biodegraded by genetically engineered microbes introduced into the soil [1].

White rot fungi (e.g. *Auricularia* sp, *Pleurotus* sp. etc.) are efficient degraders of lignocellulose [3] and other polyethylene based agro-industrial waste. Chiu and Moore [4] stated that mushrooms are efficient nutrient “recyclers”, effective in unlocking and recovering quality nutrient from recalcitrant and non-degradable waste. Lin *et al*. [5] reported that agro-industrial wastes were suitable for cultivation of *Auricularia polytricha* (An edible white rot mushroom). In China, *Auricularia* mushrooms were cultivated in corn fields such that the spent compost from the mushrooms was used as organic fertilizer for growing the corn, while the postharvest waste was used as substrate for mushroom cultivation [6].

The *Auricularia* mushrooms are very unique because they are mostly edible mushrooms, cultivated in tropical regions since their mycelia can grow at temperatures ranging from 10 to 40°C [7]. Unlike other mushroom species, cultivation of *Auricularia* is very easy and fruiting bodies are produced at shorter periods. Also, farmers do not require expensive facilities for establishment of large scale commercial mushroom farms. In addition, farmers only need to apply wastes from domestic, agricultural or agro-industrial sources for mushroom farming [8]. Hence, mushroom farming can be a perfect alibi for global waste management.

Farmers can therefore utilize agricultural wastes, which constitutes nuisance to the environment, as substrates for mushroom production [9, 10], since they are safe and devoid of heavy metals. Edosomwan *et al*. [11] extracted eggs of several helminthic parasites from commercial *Auricularia auricula* sold in Ikpoba Hill market, Benin city, Edo State, Nigeria. His findings showed that there were 53.30% of *Toxocara canis*, 13.33% of *Trichuris ovis*, and 6.67% of*Moniezia benedeni*, with heavy metals like iron (Fe) 860mg/kg, Zinc (Zn) 58mg/kg and Nickel (Ni) 1.60mg/kg present in mushrooms sold in the market, which were collected from the wild [11]. These values were higher than the recommended WHO Standards especially for Fe and Zn. Also, three genera of bacteria (*Citrobacter sp., Staphylococcus aureus*, and *Bacillus sp*.) and a fungus (*Mucor sp*.) were further isolated from the same market samples. The consumption of mushrooms with high heavy metals concentrations poses serious health risk like cancer development or heavy metal toxicity, and the presence of bacteria and helminthes eggs can result in the development of gastrointestinal infections [12].

Furthermore, the African forest is gradually undergoing destruction by man, natural disasters and wastes, alongside the germplasm of these mushrooms [13]. Therefore, there is an urgent need to rejuvenate the African environment by adopting an eco-friendly method of waste disposal and management through the development cultivation methods that will encourage rapid propagation and total conservation of endangered mushroom species. As such, a better understanding of the nutritional and physiological requirements of wild edible mushrooms will serve as a pivotal for the proposed waste management strategy that should be developed in Africa.

## 2.0 Methodology

### 2.1 Wild mushroom scavenging and sample collection

The research area was between Latitude 6.43 – 7.95°N and Longitude 2.88 – 5.24°E (Nigeria) as shown in Fig 1. The sampled region was sub-classified into States under the Southwest Province of Nigeria i.e. Oyo, Osun, Ondo, Ogun, Ekiti, and Lagos (Fig 2 and Tables 1–6). A total of 48 samples of *Auricularia* mushrooms were collected, identified, coded and geo-tagged (Tables 1–6). Laboratory analysis, spawning and cultivation of mushrooms was conducted at the National Institute of Horticulture (NIHORT), Ibadan, Nigeria and the Department of Botany, University of Ibadan, Ibadan, Nigeria.

**Fig 1.**
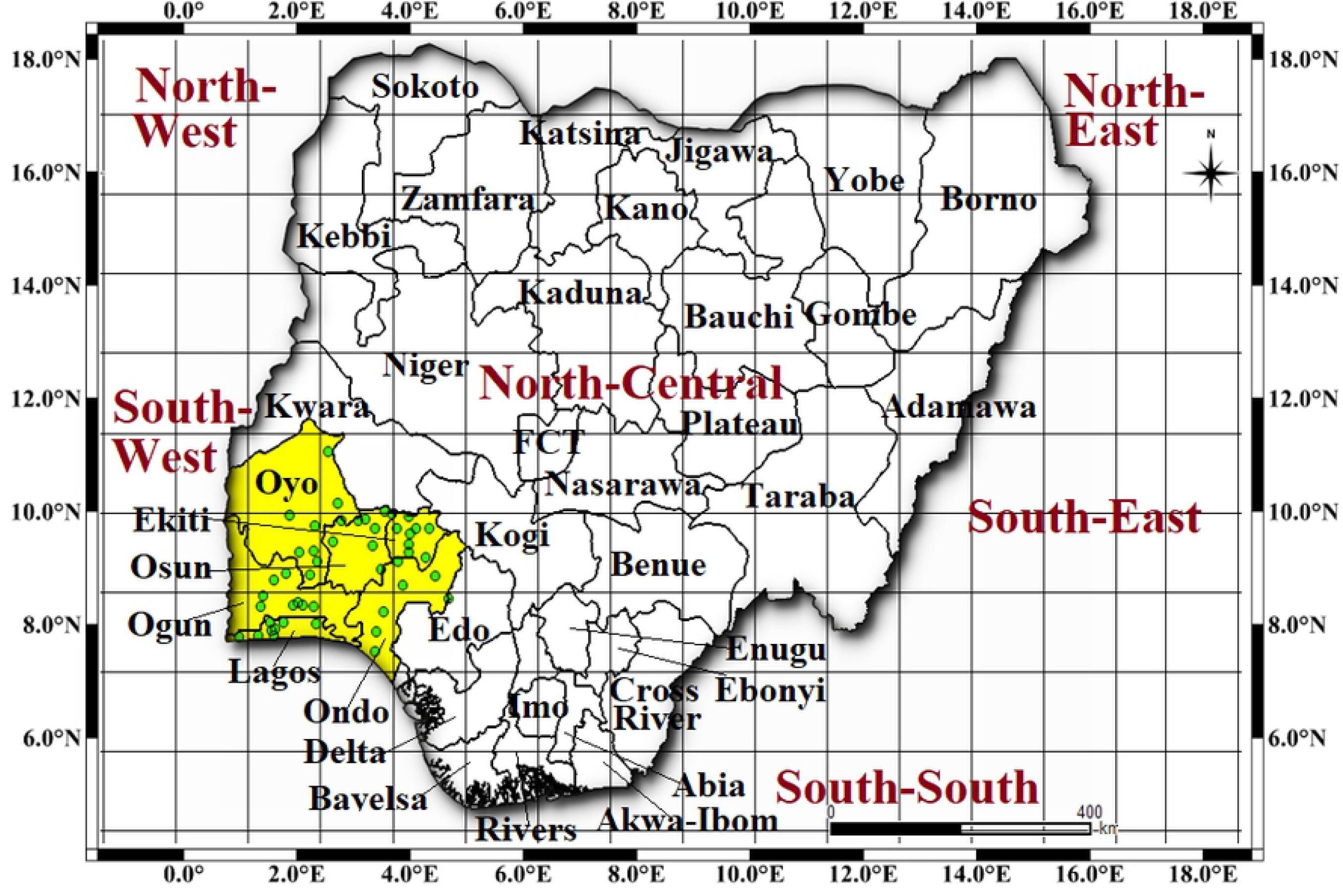
The geographical location and boundary of the research area.

**Fig 2.**
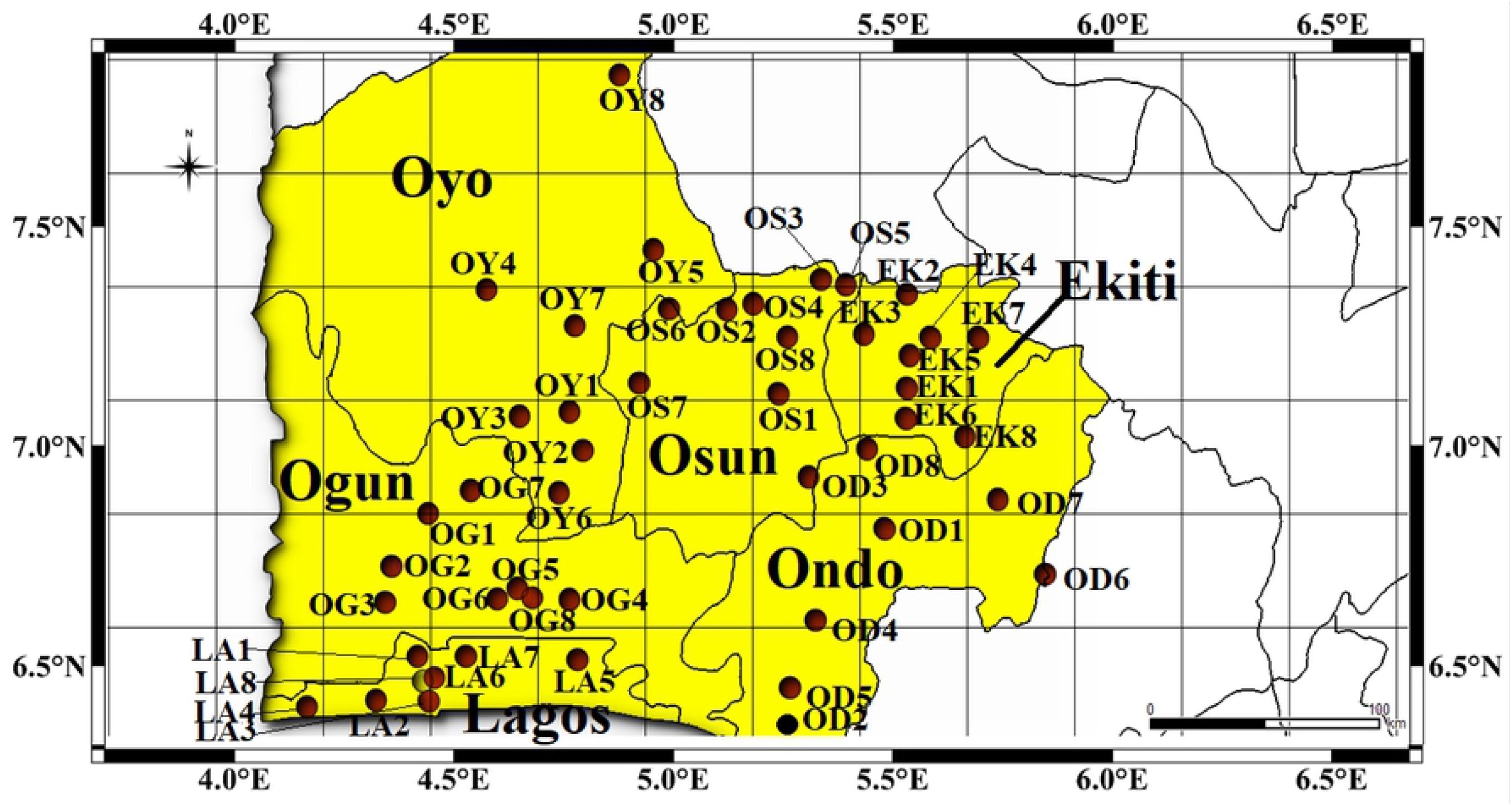
The earmarked forest locations within Southwest Province, Nigeria.

**Table 1.** Forest locations used in this research (Ogun State, Southwest-Nigeria).

| S/N | Stations | Code | Local Govt. Area | Town | State | Latitude | Longitude |
| --- | --- | --- | --- | --- | --- | --- | --- |
| 1 | 01 | OG1 | Abeokuta North | Abeokuta | Ogun | 7.1475°N | 3.3619°E |
| 2 | 02 | OG2 | Ewekoro | Itori | Ogun | 6.9530°N | 3.2181°E |
| 3 | 03 | OG3 | Ifo | Ifo | Ogun | 6.8192°N | 3.1930°E |
| 4 | 04 | OG4 | Ijebu Ode | Ijebu Ode | Ogun | 6.8300°N | 3.9165°E |
| 5 | 05 | OG5 | Ikenne | Ikenne | Ogun | 6.8717°N | 3.7105°E |
| 6 | 06 | OG6 | Shagamu | Shagamu | Ogun | 6.8322°N | 3.6319°E |
| 7 | 07 | OG7 | Odeda | Odeda | Ogun | 7.2328°N | 3.5281°E |
| 8 | 08 | OG8 | Odogbolu | Odogbolu | Ogun | 6.8365°N | 3.7689°E |

**Table 2.** The research coordinates for forest locations in Lagos State, Southwest-Nigeria.

| S/N | Stations | Code | Local Govt. Area | Town | State | Latitude | Longitude |
| --- | --- | --- | --- | --- | --- | --- | --- |
| 1 | 09 | LA1 | Agege | Ikeja | Lagos | 6.6180°N | 3.3209°E |
| 2 | 10 | LA2 | Ojo | Ojo | Lagos | 6.4579°N | 3.1580°E |
| 3 | 11 | LA3 | Apapa | Ikeja | Lagos | 6.4553°N | 3.3641°E |
| 4 | 12 | LA4 | Badagry | Badagry | Lagos | 6.4316°N | 2.8876°E |
| 5 | 13 | LA5 | Epe | Epe | Lagos | 6.6055°N | 3.9470°E |
| 6 | 14 | LA6 | Shomolu | Shomolu | Lagos | 6.5392°N | 3.3842°E |
| 7 | 15 | LA7 | Ikorodu | Ikorodu | Lagos | 6.6194°N | 3.5105°E |
| 8 | 16 | LA8 | Mushin | Ikeja | Lagos | 6.5273°N | 3.3414°E |

**Table 3.** Forest coordinates for Oyo State, Southwest-Nigeria.

| S/N | Stations | Code | Local Govt. Area | Town | State | Latitude | Longitude |
| --- | --- | --- | --- | --- | --- | --- | --- |
| 1 | 17 | OY1 | Akinyele | Moniya | Oyo | 7.5249°N | 3.9152°E |
| 2 | 18 | OY2 | Egbeda | Egbeda | Oyo | 7.3796°N | 3.9675°E |
| 3 | 19 | OY3 | Ido | Ido | Oyo | 7.5077°N | 3.7194°E |
| 4 | 20 | OY4 | Iseyin | Iseyin | Oyo | 7.9765°N | 3.5914°E |
| 5 | 21 | OY5 | Ogbomosho North | Ogbomosho | Oyo | 8.1227°N | 4.2436°E |
| 6 | 22 | OY6 | Oluyole | Idi Ayunre | Oyo | 7.2247°N | 3.8732°E |
| 7 | 23 | OY7 | Oyo | Oyo | Oyo | 7.8430°N | 3.9368°E |
| 8 | 24 | OY8 | Olorunsogo | Igbeti | Oyo | 8.7699°N | 4.1104°E |

**Table 4.** Local coordinates for forest sites in Ekiti State, Southwest-Nigeria.

| S/N | Stations | Code | Local Govt. Area | Town | State | Latitude | Longitude |
| --- | --- | --- | --- | --- | --- | --- | --- |
| 1 | 25 | EK1 | Ado Ekiti | Ado Ekiti | Ekiti | 7.6124°N | 5.2371°E |
| 2 | 26 | EK2 | Ilejemeje | Iye | Ekiti | 7.9591°N | 5.2371°E |
| 3 | 27 | EK3 | Ikole | Ikole | Ekiti | 7.7983°N | 5.5145°E |
| 4 | 28 | EK4 | Oye | Oye | Ekiti | 7.7979°N | 5.3286°E |
| 5 | 29 | EK5 | Irepodun | Igede | Ekiti | 7.7313°N | 5.2476°E |
| 6 | 30 | EK6 | Ikere | Ikere | Ekiti | 7.4991°N | 5.2319°E |
| 7 | 31 | EK7 | Ijero | Ijero Ekiti | Ekiti | 7.8120°N | 5.0677°E |
| 8 | 32 | EK8 | Emure | Emure Ekiti | Ekiti | 7.4317°N | 5.4621°E |

**Table 5.**
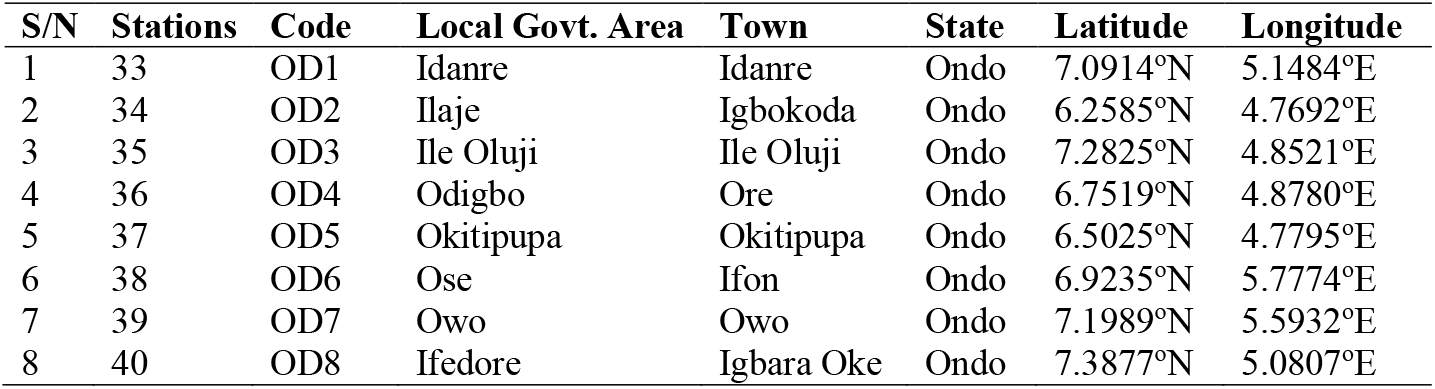
The research coordinates for Ondo State, Southwest-Nigeria.

| S/N | Stations | Code | Local Govt. Area | Town | State | Latitude | Longitude |
| --- | --- | --- | --- | --- | --- | --- | --- |
| 1 | 33 | OD1 | Idanre | Idanre | Ondo | 7.0914°N | 5.1484°E |
| 2 | 34 | OD2 | Ilaje | Igbokoda | Ondo | 6.2585°N | 4.7692°E |
| 3 | 35 | OD3 | Ile Oluji | Ile Oluji | Ondo | 7.2825°N | 4.8521°E |
| 4 | 36 | OD4 | Odigbo | Ore | Ondo | 6.7519°N | 4.8780°E |
| 5 | 37 | OD5 | Okitipupa | Okitipupa | Ondo | 6.5025°N | 4.7795°E |
| 6 | 38 | OD6 | Ose | Ifon | Ondo | 6.9235°N | 5.7774°E |
| 7 | 39 | OD7 | Owo | Owo | Ondo | 7.1989°N | 5.5932°E |
| 8 | 40 | OD8 | Ifedore | Igbara Oke | Ondo | 7.3877°N | 5.0807°E |

**Table 6.** Earmarked forest coordinates in Osun State, Southwest-Nigeria.

| S/N | Stations | Code | Local Govt. Area | Town | State | Latitude | Longitude |
| --- | --- | --- | --- | --- | --- | --- | --- |
| 1 | 41 | OS1 | Bolunduro | Ota Aiyebaju | Osun | 7.5912°N | 4.7329°E |
| 2 | 42 | OS2 | Ejigbo | Ejigbo | Osun | 7.9045°N | 4.3052°E |
| 3 | 43 | OS3 | Ifedayo | Oke Ila Orangun | Osun | 7.9946°N | 4.9974°E |
| 4 | 44 | OS4 | Ifelodun | Ikirun | Osun | 7.9227°N | 4.6347°E |
| 5 | 45 | OS5 | Ila | Ila Orangun | Osun | 8.0121°N | 4.8988°E |
| 6 | 46 | OS6 | Irepodun | Ilobu | Osun | 7.9021°N | 4.5315°E |
| 7 | 47 | OS7 | Iwo | Iwo | Osun | 7.6292°N | 4.1872°E |
| 8 | 48 | OS8 | Obokun | Ibokun | Osun | 7.8019°N | 4.7692°E |

### 2.2 Tissue culture

The method of Weber and Webster [14] was used to culture excised tissues from the pileus of the mushroom specimen. Re-hydrated Basidiocarps (*Auricularia* mushroom specimens) were washed thoroughly with 20% Teepol and sterile distilled water, after which they were sterilized in 5% sodium hypochlorite for 30s. The sterilized mushroom sections were further rinsed in three (3) successive changes of sterile distilled water and allowed to air dry for 1hour. Sterile scalpels were used to cut out fragments (about 2×2 mm^2^) from the inner surfaces of the Basidiocarp, and inoculated on freshly prepared MEA. The culture was incubated at 25±2°C for 7days. Intermittent sub-culturing was done until pure cultures were obtained.

### 2.3 Spawning on agro-industrial and market wastes

Spawn production was aided by the method of Adenipekun *et al*. [2]. Rice straw, sawdust, cotton waste and wheat bran were soaked in water for 1hr, while sorghum and maize grains were soaked for 24hrs (to remove chemical residue) after which they were drained and air dried for 4hrs (The substrates were listed in Table 7). For small scale spawn production, 500g of nutrient supplements (sorghum or maize grain) were weighed out, mixed with 1% calcium carbonate (CaCO_3_) and 1kg spawn substrate (Rice straw/Sawdust/Cotton waste and wheat bran in a ratio of 2:1, respectively). The final substrate mixture was loaded into 350cm^3^ (13 x 8 x 8) sterile bottles, covered with aluminum foil, and autoclaved at 760mmHg and 121° C for 15mins. The sterilized spawn substrate was inoculated with pure culture of *Auricularia* mushrooms and incubated at 25±2°C for three (3) weeks. For large scale spawn production, the substrate mixture was made up to 10kg and stored in heat resistant polypropylene bags (20cm x 12cm dimension). The bags were tightly secured with cotton strings and sterilized at 100°C for 4hrs. The procedure for inoculation and incubation were similar to that of the small scale spawn production method.

**Table 7.** Various substrates used in the cultivation of *Auricularia* Mushroom.

| S/N | Substrates | Agro-waste collection unit | Latitude | Longitude |
| --- | --- | --- | --- | --- |
| **<u>Nutrient Supplement</u>** |  |  |  |  |
| 1 | Wheat bran (Waste) | Bodija Market | 7.4351°N | 3.9143°E |
| 2 | Sorghum grain | Bodija Market | 7.4351°N | 3.9143°E |
| 3 | Maize grain | Bodija Market | 7.4351°N | 3.9143°E |
| 4 | Calcium carbonate | Laboratory, UI | 7.4432°N | 3.9003°E |
| **<u>Spawn Substrate</u>** |  |  |  |  |
| 5 | Sawdust (Waste) | Sawmill, Bodija | 7.4351°N | 3.9143°E |
| 6 | Cotton scraps (Waste) | Market waste, Bodija | 7.4351°N | 3.9143°E |
| 7 | Rice straw (Waste) | IITA Rice Farm | 7.3963°N | 3.9166°E |
| **<u>Growth Substrate</u>** |  |  |  |  |
| 8 | Mango (Waste wood) | Botanical Garden, UI | 7.4333°N | 3.900°E |
| 9 | Cedar (Waste wood) | Botanical Garden, UI | 7.4333°N | 3.900°E |
| 10 | Quickstick (Waste wood) | Botanical Garden, UI | 7.4333°N | 3.900°E |

### 2.4 Growth substrate prepared from garden waste

The pruned branches were cut to a standard size of 40cm using a chain saw. Holes were drilled on the logs using a manual hand drilling machine (Specification: width = 8mm, depth = 20-30mm and distance apart = 40-60mm apart). The drilled logs were covered with sterile polyethylene bags prior to inoculation to avoid contamination.

### 2.5 Mushroom Cultivation

The log technique described by Oei and Nieuwenhuijzen [15] was adopted for this research. About 1g of each spawn was aseptically inoculated into each hole on a single log at a time. The logs were incubated at 25±2°C, 760mmHg and watered regularly. After about three months, the mushrooms were ready for cropping. The number and size of the fruiting bodies after sprouting were recorded for each substrate. Yield and other agronomic parameters were also measured i.e. the number of fruiting bodies on each substrate, the height of the fruiting bodies (from the base of the stripe to the pileus), the diameter of the pileus, the fresh and dry weights, and the biomass of fruiting bodies.

***Note***: The preferred period to use log is when the leaves are just beginning to dry. Then, the sugar on the tree must have accumulated in the logs.

### 2.6 Proximate Analysis

The official methods of Association of Analytical Chemists [16] was used for the experiment.

#### 2.61 Nitrogen and Crude Protein content

1. The mushroom samples were pulverized and filtered through a 20-mesh screen and 10g of each sample was weighed out and aseptically transferred into separate Kjeldahl flasks.
2. Add 6mL each of conc. H_2_SO_4_ and catalyst (H_3_PO_4_) to each flask, and heat gently.
3. Digest the samples for 2hrs at standard temperature until complete breakdown of all organic matter was observed.

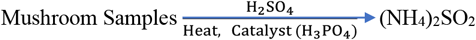
4. The digested mushroom samples were diluted with water, and Na_2_S_2_O_3_.5H_2_O was added to neutralize excess conc. H_2_SO_4_. The gas formed (NH_3_) was distilled into H_3_BO_3_ solution containing the indicators (methylene blue and methyl red).

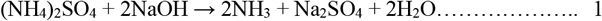

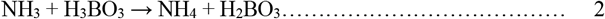
5. Titration Borate anion (proportional to the amount of nitrogen) was titrated against a standardized solution of conc. HC1.

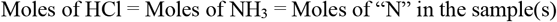

***Note***: A reagent blank should be run to subtract reagent nitrogen from the sample nitrogen

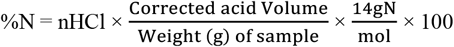

where:

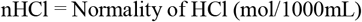

Corrected acid vol. = (ml std. acid for sample) – (ml std. acid for blank)

Atomic weight of nitrogen = 14

A factor is used to convert %N to crude protein (%).

Most proteins contain N (16%), so the conversion factor is 6.25 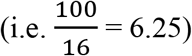.

So,

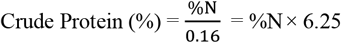

#### 2.62 Moisture content

1. Weigh 10g of the pulverized mushroom samples into separate pre-weighed crucibles (mo).
2. The crucibles containing the samples (m_1_) were heated at a steady temperature (105°C) and constant pressure (760mmHg) in a well ventilated hot air oven.
3. The weight of the loaded crucibles was measured at hourly intervals until similar values were obtained between successive measurements (m_2_).
4. The loss in weight was reported as the moisture content and calculated by the formula:

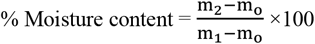 Since,

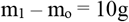 Therefore,

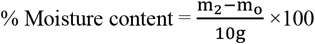 Where,

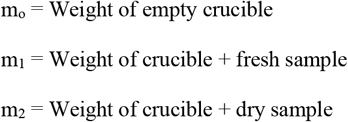

#### 2.63 Crude fat content

1. Weigh 2g of dried pulverized samples into different extraction thimbles with porosity that can allow a rapid flow of (C_2_H_5_)_2_O and cover the thimbles with glass wool
2. Weigh clean boiling flasks and add anhydrous (C_2_H_5_)_2_O.
3. Add small pieces of Na metals to inhibit the oxidation of hydrogen
4. Assemble boiling flask, Soxhlet flask, and condenser accordingly.
5. Extract fats in a Soxhlet extractor at a rate of 5 or 6 drops per second by condensation for about 4hrs, or for 16hrs at a rate of 2 or 3 drops per second by heating solvent in boiling flask.
6. Dry boiling flask with extracted fat in a hot air oven at 100*°*C for 30min
7. Cool in a desiccator and weigh.

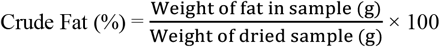

#### 2.64 Crude fiber content

1. Weigh 2g of defatted mushroom sample into 500ml conical flask
2. Add 200ml of 1.25% (0.255M) H_2_SO_4_ and heat at 100°C for 30mins, maintaining a constant volume (using reflux condenser).
3. Rotate the flask every 10mins to enhance homogenization.
4. Add 20ml of 28% KOH to the mixture while still heating at 100°C for another 30mins.
5. Add a mixture of 1% H_2_SO_4_ and 1.25% (0.313M) NaOH solution afterwards
6. Filter and dry the content of the flask in an electric oven at 130° C to a constant weight for 2hrs.
7. The filtrate was then left in a desiccator at 25±2°C for 30mins to dry further
8. Weigh the filtrates before transferring into a crucible
9. Place the pre-weighed crucibles in a muffle furnace at 550°C for complete ashing.
10. The ash obtained was left to cool down in a desiccator and re-weighed.
11. The crude fiber content was determined by using the formula.

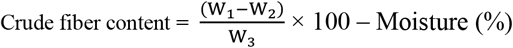

Where,

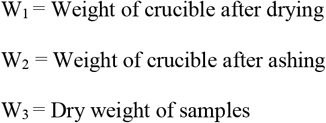

#### 2.65 Ash content

1. Weigh 5-10g of the mushroom sample into a tared crucible.
2. Place crucibles in muffle furnace and heat for 12-18hrs (or overnight) at about 550*°*C.
3. Transfer crucibles to a desiccator with a porcelain plate and desiccant.
4. Cover crucibles, close desiccator, and allow crucibles to cool prior to weighing.
5. The ash content is calculated as follows

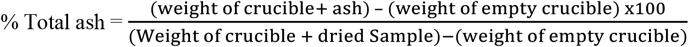

#### 2.66 Carbohydrate content

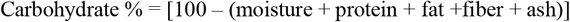

### 2.7 Mineral Analysis

The official methods of Association of Analytical Chemists [16] was used for the experiment.

#### 2.71 Phosphorus (P)

1. Transfer the solution containing the ash from the incinerated mushroom samples to a 250mL beaker.
2. Add NH_4_OH in excess. Dissolve the precipitate formed with a few drops of HNO3.
3. Add 70mL of MoO4^-2^ solution for every decigram of P_4_O_10_ present.
4. Digest at about 65°C. for 1hr until there is complete precipitation of P_4_O_10_.
5. Filter, and wash with cold water.
6. Dissolve the precipitate with NH_4_OH (1:1) and hot water. Equate the volume to 100mL.
7. Neutralize with HC1, using litmus paper or bromothymol blue as indicator.
8. Titrate slowly, at about 1 drop per second, stirring vigorously, 15 mL of the magnesia mixture for each decigram of P_4_O_10_ present.
9. After 15 minutes, add 12 mL of NH_4_OH. Allow to stand for 2hrs, filter and wash the precipitate with NH_4_OH (1:9) until the washings are practically free of chlorides.
10. Dry in a dessicator, burn at a low heat, and then ignite in an electric furnace at 950-1000°C.
11. Cool in a desiccator and weigh as Mg_2_P_2_O_7_
12. Calculate the result as % P_4_O_10_.

#### 2.72 Potassium (K)

The Gravimetric procedure was used to determine the level of potassium in the samples.

1. Prepare an aqueous solution containing 1g of the Na_3_Co(NO_2_)_6_ in 5mL solution. Filter before use.
2. The aliquot or sample for analysis should contain between 2 and 15 mg. of potassium in a neutral aqueous solution of 10 mL volume.
3. Add 1mL of 1N HNO3 and 5mL of Na_3_Co(NO_2_)_6_ solution, mix, and allow to stand for 2hrs.
4. Filter in a porous-bottomed porcelain crucible, the tare weight of which is known, using 0.01N HNO3 in a wash bottle to make the transfer.
5. Wash 10 times with 2mL portions of the dilute HNO3 and 5 times with 2mL portions of 95% alcohol.
6. Dry for 1hr at 110°C. Cool in a desiccator, and weigh.
7. The composition of the precipitate can be represented by the formula

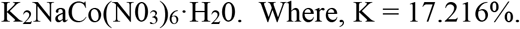

#### 2.73 Magnesium (Mg)

1. The samples were digested using a mixture of concentrated HNO3, H_2_SO_4_ and HC10_4_ (10:0:5:2, v/v).
2. Thereafter, the contents were analyzed using the atomic absorption spectrophotometer (GBC 904AA; Germany).

#### 2.74 Calcium (Ca)

1. Weigh 10g of the ashed samples into a porcelain dish, about 60 mm in diameter and having a capacity of 35mL
2. Dissolve by adding 2mL of hydrochloric acid with the aid of a pipette. Cover the dish with a watch glass, heat for 5 minutes on a steam bath, wash off the watch glass, and filter.
3. Transfer the filtrate to a 100mL volumetric flask. Cool and dilute to 50mL volume.
4. Add 8 to 10 drops of bromocresol green indicator solution and 20% C_2_H_3_NaO_2_. Cover with a watch glass and heat to boiling.
5. Add slowly 3% C_2_H_2_O_4_ solution, a drop every 3-5s, until it turns green or the pH is reduced to 4.4 or 4.6.
6. Boil for 1-2mins and allow to stand overnight.
7. Filter the supernatant solution through a sintered-glass filter
8. Titrate at 70-90°C with 0.05 *N*KMnO_4_ solution until a slight pink color is obtained.
9. Run a blank determination and make a correction. 1mL of 0.05 *N* KMnO4 solution is equivalent to 1 mg. of calcium.

#### 2.75 Sodium (Na)

1. Transfer 10-15mL aliquot of sample solution in a 50mL beaker.
2. Add an excess of powdered ZnCO_3_, cover the beaker, and let stand at 25±2°C for 24hrs.
3. Evaporate to dryness, add 10 mL of H_2_O and conc. HC1 to dissolve the salts. Then add the zinc carbonate.
4. Filter through quantitative paper in a small funnel and wash thoroughly with cold water 5 or 6 times with small portions of water.
5. Decant the filtrate and washings in a small beaker and evaporate to 1 or 2mL Add 100mL C_4_H_6_O_6_UZn reagent and stir vigorously for 1hr and allow to stand overnight.
6. Collect the filtrates through a weighed Gooch crucible, wash with 95% alcohol saturated with the NaZn(UO_2_)_3_(C_2_H_3_O_2_)_9_ salt and then with ether saturated with the same salt.
7. Dry in air and weigh. The salt has the formula: NaZn(UO_2_)3(C_2_H_3_O_2_)9·6H_2_O.

### 2.8 Data Analysis

The data obtained were analyzed using the Statistical Analysis Software [17] Version 9.4, while the homogeneity of means was ascertained by Tukey’s test (P<0.05).

## 3.0 Results

The research conducted was aimed at developing easier and faster methods of cultivating estranged mushroom species that are of paramount importance to man and the environment. The research took into consideration various growth stages for mushroom development in order to determine the best combination of nutrient substrate at every phase.

### 3.1 Spawning

Postharvest waste from the IITA rice field reduced the spawning periods of mushroom samples from the wild forest of Ekiti (EK1-8 = 23days), Lagos (LA1-8 = 21days), Ogun (OG1-8 = 20days) and Oyo (OY1-8 = 22.13days) as stated in Table 8. Also, cotton waste from Bodija market was beneficial in reducing the spawn period of mushroom samples collected from Ondo (OD1-8 = 23days) and Osun (OS1-8 = 20days). The quality of spawn substrate was rated as Rice straw>Cotton waste>Saw dust (Table 8).

**Table 8.**
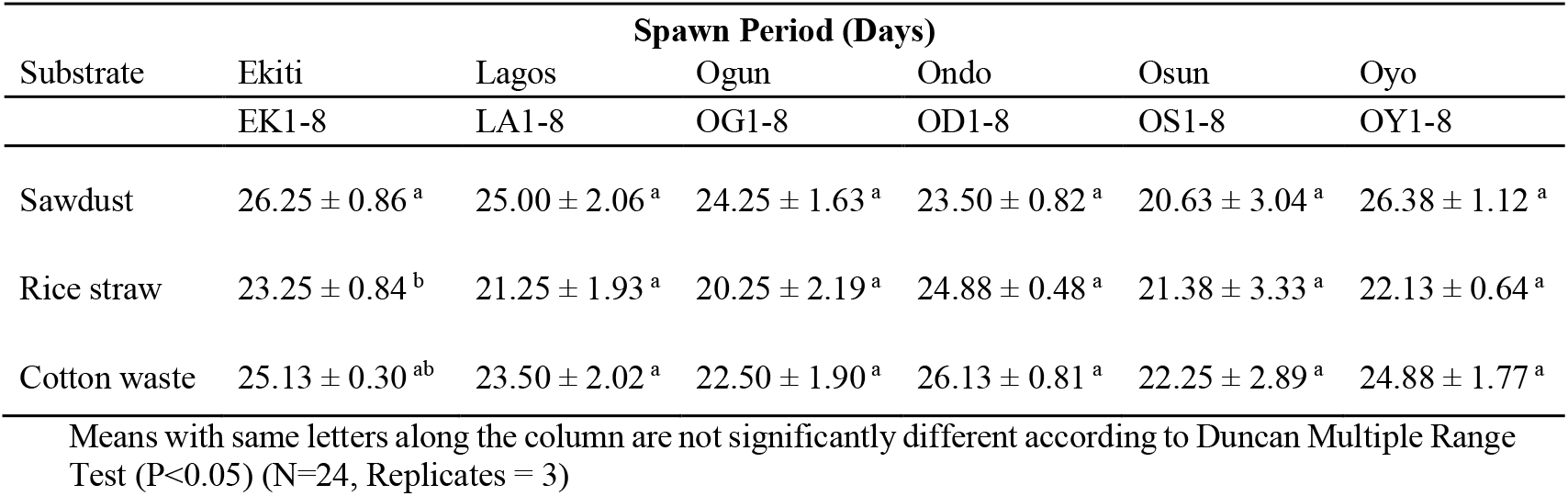
Spawn production from different wastes.

| Substrate | Spawn Period (Days) |  |  |  |  |  |
| --- | --- | --- | --- | --- | --- | --- |
|  | Ekiti | Lagos | Ogun | Ondo | Osun | Oyo |
|  | EK1-8 | LA1-8 | OG1-8 | OD1-8 | OS1-8 | OY1-8 |
| Sawdust | 26.25 ± 0.86<sup>a</sup> | 25.00 ± 2.06<sup>a</sup> | 24.25 ± 1.63<sup>a</sup> | 23.50 ± 0.82<sup>a</sup> | 20.63 ± 3.04<sup>a</sup> | 26.38 ± 1.12<sup>a</sup> |
| Rice straw | 23.25 ± 0.84<sup>b</sup> | 21.25 ± 1.93<sup>a</sup> | 20.25 ± 2.19<sup>a</sup> | 24.88 ± 0.48<sup>a</sup> | 21.38 ± 3.33<sup>a</sup> | 22.13 ± 0.64<sup>a</sup> |
| Cotton waste | 25.13 ± 0.30<sup>ab</sup> | 23.50 ± 2.02<sup>a</sup> | 22.50 ± 1.90<sup>a</sup> | 26.13 ± 0.81<sup>a</sup> | 22.25 ± 2.89<sup>a</sup> | 24.88 ± 1.77<sup>a</sup> |

Means with same letters along the column are not significantly different according to Duncan Multiple Range Test (P<0.05) (N=24, Replicates = 3)

### 3.2 Pin head emergence

The fastest time for pin head emergence of most of the cultivated *Auricularia* mushrooms (83.3%) collected at different locations from the wild was observed on *G. sepium* substrate i.e. EK1-8 (26days), LA1-8 (25days), OG1-8 (24days), OS1-8 (25days) and OY1-8 (29days), respectively (Table 9). Pin head formation was best supported by *Mangifera indica* substrate for mushroom samples collected from the tropical forest in Ondo State (OD1-8 = 26days).

**Table 9.** Duration for pin head emergence.

| Substrate | Pin Head Emergence (Days) |  |  |  |  |  |
| --- | --- | --- | --- | --- | --- | --- |
|  | Ekiti | Lagos | Ogun | Ondo | Osun | Oyo |
|  | EK1-8 | LA1-8 | OG1-8 | OD1-8 | OS1-8 | OY1-8 |
| *M. indica* | 29.13 ± 0.69<sup>a</sup> | 29.00 ± 1.50<sup>a</sup> | 28.75 ± 0.80<sup>a</sup> | 26.50 ± 0.71<sup>b</sup> | 26.50 ± 2.19<sup>a</sup> | 32.13 ± 1.30<sup>a</sup> |
| *G. sepium* | 26.50 ± 0.87<sup>b</sup> | 25.50 ± 1.71<sup>a</sup> | 24.75 ± 1.54<sup>a</sup> | 27.75 ± 0.31<sup>ab</sup> | 25.75 ± 2.55<sup>a</sup> | 29.13 ± 1.14<sup>a</sup> |
| *C. odorata* | 28.25 ± 0.53<sup>ab</sup> | 27.13 ± 1.54<sup>a</sup> | 26.75 ± 1.19<sup>a</sup> | 29.00 ± 0.73<sup>a</sup> | 26.50 ± 2.12<sup>a</sup> | 30.63 ± 1.67<sup>a</sup> |

Means with same letters along the column are not significantly different according to Duncan Multiple Range Test (P<0.05) (N=24, Replicates = 3)

### 3.3 Formation of mushroom fruiting body

The best substrate for fruiting of *Auricularia* mushrooms was ranked as follow: *G. sepium*>*C. odorata>M. indica* (Table 10). *G. sepium* supported the early fruiting body formation of cultivated mushrooms (83.3%) collected at different strategic locations in the wild i.e. Ekiti (28days), Lagos (27days), Ogun (27days), Osun (28days) and Oyo (31days), respectively (Table 10). *C. odorata* only supported the optimum duration for fruiting body formation in wild mushroom specimens (16.7%) collected from Ondo (29days) as shown in Table 10.

**Table 10.** Duration for formation of fruiting bodies.

| Substrate | Mushroom Fruiting Body Formation (Days) |  |  |  |  |  |
| --- | --- | --- | --- | --- | --- | --- |
|  | Ekiti | Lagos | Ogun | Ondo | Osun | Oyo |
|  | EK1-8 | LA1-8 | OG1-8 | OD1-8 | OS1-8 | OY1-8 |
| *M. indica* | 31.13 ± 0.69<sup>a</sup> | 30.88 ± 1.48<sup>a</sup> | 30.75 ± 0.80<sup>a</sup> | 29.00 ± 0.66<sup>a</sup> | 28.75 ± 2.17<sup>a</sup> | 34.00 ± 1.23<sup>a</sup> |
| *G. sepium* | 28.63 ± 0.84<sup>b</sup> | 27.38 ± 1.68<sup>a</sup> | 27.13 ± 1.33<sup>a</sup> | 30.13 ± 0.40<sup>a</sup> | 28.13 ± 2.42<sup>a</sup> | 31.50 ± 1.00<sup>a</sup> |
| *C. odorata* | 30.38 ± 0.46<sup>ab</sup> | 29.38 ± 1.57<sup>a</sup> | 28.75 ± 1.15<sup>a</sup> | 30.75 ± 0.65<sup>a</sup> | 28.63 ± 1.97<sup>a</sup> | 32.13 ± 1.62<sup>a</sup> |

Means with same letters along the column are not significantly different according to Duncan Multiple Range Test (P<0.05) (N=24, Replicates = 3)

### 3.4 Mushroom harvesting periods

The maturation period for the cultivated *Auricularia* mushrooms was reduced when grown on *Cedrela odorata* substrate i.e. EK1-8 (2days), while LA1-8, OG1-8, OD1-8 and OY1-8 was 3days (Table 11). Other substrates too had positive influence in the reduction of harvesting period too (Table 11). The matured fruiting bodies of the cultivated *Auricularia* mushrooms on waste woods (*Mangifera indica*, *Gliricidium sepium* and *Cedrela odorata*).

**Table 11.** Harvesting periods for each mushroom.

| Substrate | Harvesting Period (Days after fruiting body formation) |  |  |  |  |  |
| --- | --- | --- | --- | --- | --- | --- |
|  | Ekiti | Lagos | Ogun | Ondo | Osun | Oyo |
|  | EK1-8 | LA1-8 | OG1-8 | OD1-8 | OS1-8 | OY1-8 |
| *M. indica* | 3.00 ± 0.38<sup>a</sup> | 3.25 ± 0.45<sup>a</sup> | 3.38 ± 0.32<sup>a</sup> | 3.50 ± 0.42<sup>a</sup> | 3.13 ± 0.40<sup>a</sup> | 4.00 ± 0.50<sup>a</sup> |
| *G. sepium* | 2.50 ± 0.27<sup>a</sup> | 3.50 ± 0.46<sup>a</sup> | 3.38 ± 0.42<sup>a</sup> | 3.25 ± 0.45<sup>a</sup> | 3.13 ± 0.30<sup>a</sup> | 3.25 ± 0.25<sup>a</sup> |
| *C. odorata* | 2.50 ± 0.27<sup>a</sup> | 3.13 ± 0.30<sup>a</sup> | 3.13 ± 0.35<sup>a</sup> | 3.13 ± 0.23<sup>a</sup> | 3.50 ± 0.33<sup>a</sup> | 3.00 ± 0.38<sup>a</sup> |

Means with same letters along the column are not significantly different according to Duncan Multiple Range Test (P<0.05) (N=24, Replicates = 3)

### 3.5 Measurement of the radial diameter of the pileus

The size of the wild *Auricularia* mushrooms from Ekiti and Oyo States were greatly improved after cultivation on waste wood from *Mangifera indica* (EK1-8: 4.59 and OY1-8: 4.51cm, respectively) when cultivated on (Table 12). Agro-waste from *Gliricidium sepium* also supported the optimal improvement in size of the cultivated mushrooms collected from the wild tropical forest of Ogun (OG1-8 = 4.58cm), Ondo (OD1-8 = 4.40cm) and Osun (OS1-8 = 4.34cm) States, respectively (Table 12), while *C. odorata* just had an optimum effect on the size of *Auricularia* mushrooms collected in Lagos State only (LA1-8 = 4.39cm).

**Table 12.**
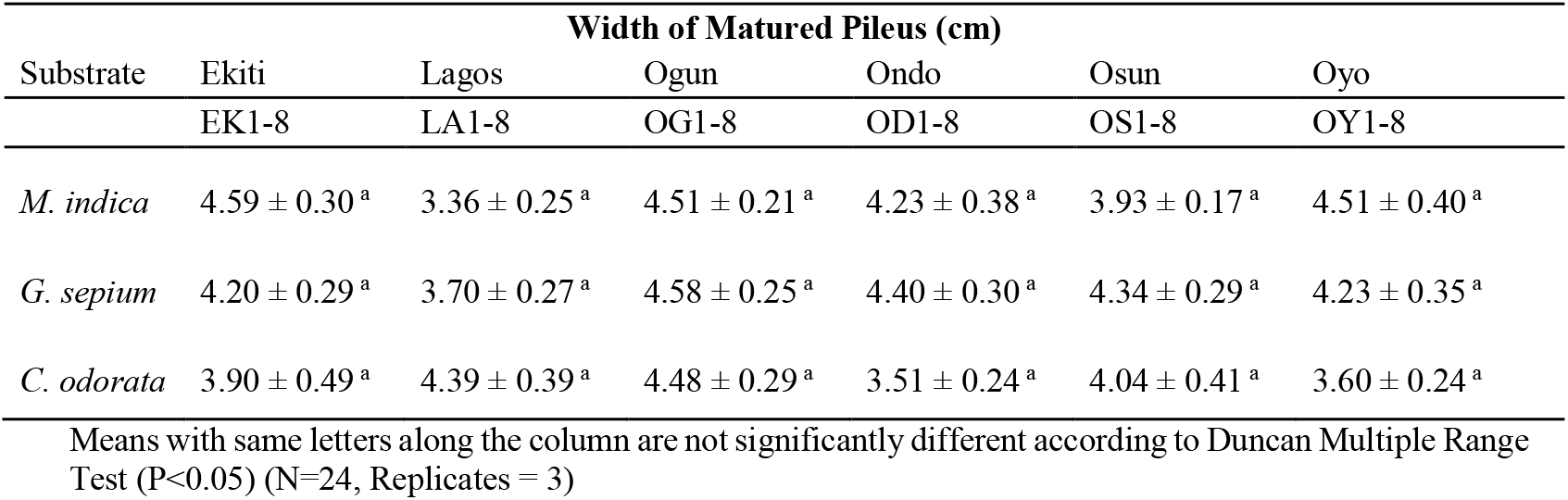
Measurement of pileus girth.

| Substrate | Width of Matured Pileus (cm) |  |  |  |  |  |
| --- | --- | --- | --- | --- | --- | --- |
|  | Ekiti | Lagos | Ogun | Ondo | Osun | Oyo |
|  | EK1-8 | LA1-8 | OG1-8 | OD1-8 | OS1-8 | OY1-8 |
| *M. indica* | $4.59 \pm 0.30^a$ | $3.36 \pm 0.25^a$ | $4.51 \pm 0.21^a$ | $4.23 \pm 0.38^a$ | $3.93 \pm 0.17^a$ | $4.51 \pm 0.40^a$ |
| *G. sepium* | $4.20 \pm 0.29^a$ | $3.70 \pm 0.27^a$ | $4.58 \pm 0.25^a$ | $4.40 \pm 0.30^a$ | $4.34 \pm 0.29^a$ | $4.23 \pm 0.35^a$ |
| *C. odorata* | $3.90 \pm 0.49^a$ | $4.39 \pm 0.39^a$ | $4.48 \pm 0.29^a$ | $3.51 \pm 0.24^a$ | $4.04 \pm 0.41^a$ | $3.60 \pm 0.24^a$ |

Means with same letters along the column are not significantly different according to Duncan Multiple Range Test ( $P < 0.05$ ) (N=24, Replicates = 3)

### 3.6 Dry matter determination

For samples collected from the wild forest region of Ekiti, Ogun, Ondo and Osun State, there was no significant difference (P<0.05) in the dry matter composition of all the mushrooms cultivated on different substrates, but the highest weight recorded was 3.02g on *G. sepium* and *C. odorata* (Ekiti), 3.04g on *C. odorata* (Ogun) and*M. indica* (Osun), 3.05g on *G. sepium* (Ondo), respectively (Table 13). The best dry mass recorded for all harvested *Auricularia* mushroom fruiting bodies from Lagos and Oyo origin was supported by*M. indica* (3.04g) and *C. odorata* (3.02g) substrates, respectively.

**Table 13.** Dry weight measurement.

| Substrate | Dry Weight (g) |  |  |  |  |  |
| --- | --- | --- | --- | --- | --- | --- |
|  | Ekiti | Lagos | Ogun | Ondo | Osun | Oyo |
|  | Ek1-8 | LA1-8 | OG1-8 | OD1-8 | OS1-8 | OY1-8 |
| *M. indica* | $2.94 \pm 0.06^a$ | $3.04 \pm 0.03^a$ | $2.95 \pm 0.06^a$ | $2.99 \pm 0.03^a$ | $3.04 \pm 0.04^a$ | $2.84 \pm 0.05^b$ |
| *G. sepium* | $3.02 \pm 0.03^a$ | $2.84 \pm 0.07^b$ | $2.98 \pm 0.05^a$ | $3.05 \pm 0.04^a$ | $2.97 \pm 0.05^a$ | $2.99 \pm 0.05^{ab}$ |
| *C. odorata* | $3.02 \pm 0.04^a$ | $2.99 \pm 0.04^{ab}$ | $3.04 \pm 0.05^a$ | $3.01 \pm 0.03^a$ | $2.97 \pm 0.07^a$ | $3.02 \pm 0.05^a$ |

Means with same letters along the column are not significantly different according to Duncan Multiple Range Test ( $P < 0.05$ ) (N=24, Replicates = 3)

### 3.7 Average yield determination

There was no significant difference (P<0.05) in the turn over (yield) of the harvested *Auricularia* mushrooms (Table 14). Though, the highest yield (In terms of value) was from samples EK1-8 (8.60g) and LA1-8 (9.52g), grown on *C. odorata* substrate. Samples from Ogun and Oyo States had better yield on pruned branches of *G. sepium* too (OG1-8 = 9.15 and OY1-8 = 9.71g), while those from Ondo and Osun States had average yields of 8.84 and 9.35g, respectively on *Mangifera indica* substrate (Table 14).

**Table 14.** Average yield evaluation.

| Substrate | Average Yield (g) |  |  |  |  |  |
| --- | --- | --- | --- | --- | --- | --- |
|  | Ekiti | Lagos | Ogun | Ondo | Osun | Oyo |
|  | EK1-8 | LA1-8 | OG1-8 | OD1-8 | OS1-8 | OY1-8 |
| *M. indica* | $8.58 \pm 0.52^a$ | $8.89 \pm 0.24^a$ | $8.99 \pm 0.58^a$ | $8.84 \pm 0.55^a$ | $9.35 \pm 0.48^a$ | $9.65 \pm 0.48^a$ |
| *G. sepium* | $8.03 \pm 0.29^a$ | $9.38 \pm 0.50^a$ | $9.15 \pm 0.53^a$ | $8.79 \pm 0.54^a$ | $9.04 \pm 0.61^a$ | $9.71 \pm 0.49^a$ |
| *C. odorata* | $8.60 \pm 0.59^a$ | $9.52 \pm 0.52^a$ | $8.22 \pm 0.31^a$ | $8.28 \pm 0.20^a$ | $8.72 \pm 0.44^a$ | $9.70 \pm 0.60^a$ |

Means with same letters along the column are not significantly different according to Duncan Multiple Range Test ( $P < 0.05$ ) (N=24, Replicates = 3)

### 3.8 Summary of growth requirements for mushroom farming

The best substrate for spawn production, pin head emergence and production of mushroom fruiting bodies was *Gliricidium sepium* with a short time frame of 22days for spawn production, 26days for pin head emergence, and 28days for the *Auricularia* mushrooms to produce fruiting bodies, on the average (Table 15). The best substrate that supports early harvesting of mushrooms was *C. odorata* with an average value of 3days. The best substrate for yield improvement, enhanced size of mushrooms produced and increase in dry matter acquisition were *M. indica* (9.05g), *G. sepium* (4.24g) and *C. odorata* (3.01g), respectively (Table 15).

**Table 15.**
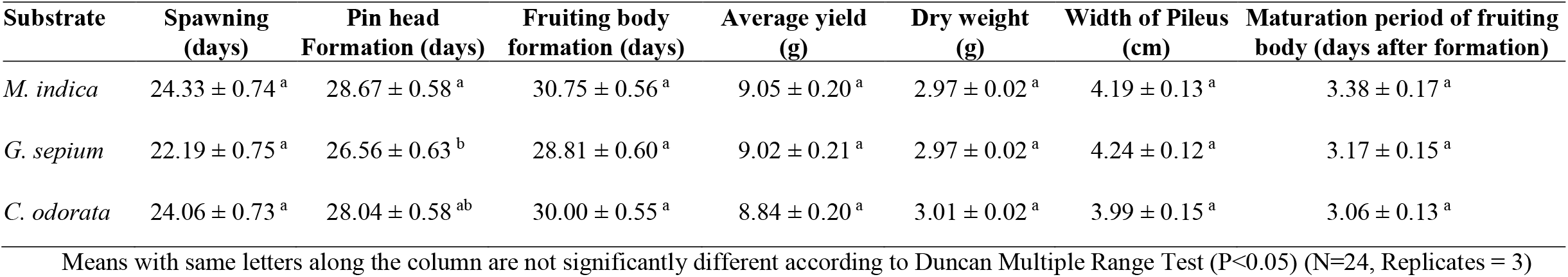
Summary of growth requirements for mushroom cultivation.

### 3.9 Mineral composition of the harvested mushrooms

Samples LA1-8 had the best Nitrogen (N), Sodium (Na) and Calcium (Ca) contents (13.59, 70.49 and 61.90mg/100g, respectively), while sample OG1-8 had the best Phosphorus composition (39.2mg/100g). The best Potassium composition was found in sample OS1-8 (1511.63mg/100g) as presented in Table 16.

**Table 16.**
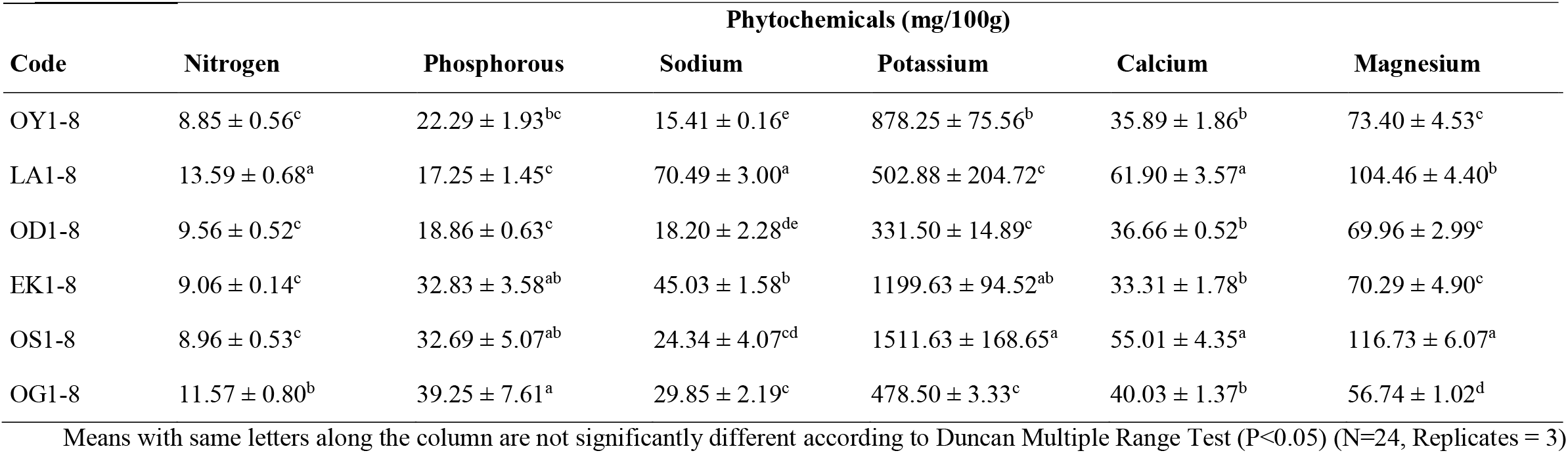
Mineral composition of *Auricularia* sp.

### 3.10 Proximate composition of the mushrooms

There was no significant difference (P<0.05) in the protein and ash contents of all the *Auricularia* mushrooms cultivated in the present study irrespective of the substrates used to grow them and the location where they were collected from (Table 17). The highest fat content was obtained from samples EK1-8 (6.60%), while the highest crude fibre content was found in samples OS1-8 (25.13%), and carbohydrate composition was more in OY1-8 (53.95%), LA1-8 (53.38%) and OG1-8 (54.23%), as shown in Table 17.

**Table 17.** Proximate analysis of the harvested *Auricularia* sp.

| Code | % Protein | % Ash | % Moisture | % Fat | % Crude Fibre | % CHO |
| --- | --- | --- | --- | --- | --- | --- |
| OY1-8 | 6.26 ± 0.41<sup>a</sup> | 4.40 ± 0.63<sup>a</sup> | 12.49 ± 0.59<sup>b</sup> | 3.60 ± 0.16<sup>bc</sup> | 19.31 ± 0.35<sup>c</sup> | 53.95 ± 1.08<sup>a</sup> |
| LA1-8 | 6.98 ± 0.81<sup>a</sup> | 4.29 ± 0.52<sup>a</sup> | 14.10 ± 0.32<sup>b</sup> | 2.76 ± 0.08<sup>c</sup> | 18.49 ± 0.37<sup>c</sup> | 53.38 ± 0.89<sup>a</sup> |
| OD1-8 | 6.24 ± 0.10<sup>a</sup> | 4.83 ± 0.42<sup>a</sup> | 16.22 ± 0.65<sup>a</sup> | 3.41 ± 0.15<sup>c</sup> | 20.51 ± 0.42<sup>bc</sup> | 48.80 ± 0.94<sup>bc</sup> |
| EK1-8 | 6.70 ± 0.17<sup>a</sup> | 3.87 ± 0.72<sup>a</sup> | 12.82 ± 0.68<sup>b</sup> | 6.60 ± 0.14<sup>a</sup> | 18.66 ± 0.28<sup>c</sup> | 51.35 ± 0.88<sup>ab</sup> |
| OS1-8 | 6.55 ± 0.16<sup>a</sup> | 4.72 ± 0.43<sup>a</sup> | 13.01 ± 1.28<sup>b</sup> | 4.41 ± 0.32<sup>b</sup> | 25.13 ± 2.48<sup>a</sup> | 46.19 ± 2.48<sup>c</sup> |
| OG1-8 | 5.31 ± 0.14<sup>a</sup> | 3.14 ± 0.12<sup>a</sup> | 10.43 ± 0.19<sup>c</sup> | 4.36 ± 0.59<sup>b</sup> | 22.54 ± 0.49<sup>ab</sup> | 54.23 ± 1.03<sup>a</sup> |

Means with same letters along the column are not significantly different according to Duncan Multiple Range Test (P<0.05) (N=24, Replicates = 3)

## 4.0 Discussion

The spawning period for the domesticated *Auricularia* mushrooms was reduced by the substrates used as growth media. Therefore, in a shorter period of time mllions of pure disease-free and non-toxic mushroom spawns can be produced in commercial quantity to meet up with the rising global demand for edible mushrooms. Also, the large scale production aided by the outcome of this research can help propel the continuous involvement of mushrooms in areas of bioremediation, biotechnology, biodetoxification, mushroom husbandary and biodiversity, genetics and medicine etc. without over stressing or allienating the mushroom gene pool to a point of extinction. This research was in agreement with the findings of Thiribhuvanamala *et al.*, [18], who reported that paddy straw, mixed saw dust and wheat bran substrates resulted in early spawning of the cultivated mushrooms with uniform mycelia growth of *A. polytricha* with bio efficiency of 46.4%.

*Gliricidium sepium* substrate had tremendous effects in the formation of mushroom pin heads and mushroom fruiting bodies in the current research conducted. This can be attributed to the specificity in the nutrient composition of the wood. Although, not scientifically proven, the wood of *G. sepium* may contain some chemical substances that can help stimulate spore formation, pin head emergence and fruiting body production in *Auricularia* mushrooms from Southwest, Nigeria. However, the rapid production of mushroom pin heads is a welcome development for producers of edible mushrooms in Africa and around the world as pin head formation is a precursor of fruiting body production, which of course is the edible aspect of the cultivated mushrooms. This was in line with the findings of Rajput and Rao [19], who noted that dead and moist part of the bark of *Mangifera indica* L supports germination of fungal spores and the growth of fungal hyphae of saprophytic fungi like *Auricularia* mushrooms. Also, Veeralakshmi *et al*. [20], noted that a combination of paddy straw + wheat bran (3:1) ratio recorded minimum days for spawn run (21days), pin head formation (31days) and first harvest (35days).

The harvesting period for these mushrooms was greatly reduced in the germplasm collected from wild mushroom specimens cultvated on *Cedrela odorata* substrate. This is an indication that several substrates shoud be used at different growth stage in other to obtain optimum results and maximize production output. This was in line with the observation of Onyango *et al*. [21], who reported that organic substrates and nutrient supplements vary proportionally in their suitability for wood ear mushroom (*Auricularia*) cultivation. Also, there was no significant difference in the total turn over (Yield) of the cultivated mushrooms regardless of the substrates used or nutrient supplements applied. Therefore, a more cheaper and readily available nutrient supplement can be used to grow *Auricularia* mushrooms without unecessary anxiety regarding the tone down of the final outcome. This was in accordance with the observations made by Adenipekun *et al.*, [2], that substrates chosen and used in mushroom cultivation are numerous and can include both field-based residues (oil palm frond, corn husk, rice straw and wheat straw etc.) and also processing based-residues (sugarcane bagasse, brewers/spent grains, rice bran and palm pressed fiber etc.).

There was negligible difference in the dry matter composition of all the cultivated mushrooms investigated in this research irrespective of the source of germplasm, but the sizes of the mushroom was influenced by the substrate used. Therefore, it is advicaeble to apply a substrate that can produce greater physical appearance of the muchrooms because mushroom lovers are mostly attracted by the size or robustness of the mushrooms. This observation was in concordance with the research conducted by Chang *et al*. [22] who noticed substantial increase in the yield of fruiting bodies per unit weight by addition of supplements to wheat bran substrate in oyster mushroom cultivation. Wang [23] also reported that supplementation of fruiting substrate resulted in a significant increase of oyster mushroom yield, improve the production, quality, flavour and also shelf life of cultivated mushrooms.

*Auricularia* mushroom germplasm from the wild forest of Lagos State cultivated in the course of this research, had greater propensity for production of beneficial mineral nutrients than those from Osun, Ogun, Ondo, Ekiti and Oyo States. This was in line with the findings of Chang and Quimio [24], who stated that most fungi are able to synthesize their own vitamins. There was no significant difference in the protein and ash contents of all the *Auricularia* mushrooms cultivated in the present study irrespective of the substrates used to grow them and the location where they were collected from. The highest fat content was obtained from mushrooms samples from Ekiti State, while the highest crude fibre content was from Osun samples, and carbohydrate composition was more in mushroom samples from Oyo, Lagos and Ogun States. This observation was synonymous to that made by Okhuoya and Ayodele [25], who noted that fresh mushrooms contain relatively large amounts of carbohydrate and fibre ranging from 51 to 88% and 4 to 20% (dry weight), respectively, for the major cultivated species.

## 5.0 Conclusion

The generation of profitable income through wastes disposal and waste wood management is a more lucrative aspect for rapid economic growth that has not yet been fully exploited globally. Cultivation of mushrooms on industrial wastes, waste woods, agricultural wastes or wastelands can foster rapid biological cleaning of the environment, create more arable farmlands for the cultivation of other economically important food crops, increase the option of nutrient availabity to man and animals, promote wealth generation from waste, and finally, reduce risk of mycotoxicosis through the consumption of poisonous mushrooms reported globally. The findings from this research is a pointer to the endless possibilities of wealth acquisition from wastes.

## Ethical Statement

This is to confirm that:

Prof. Clementina O. Adenipekun, Dr. V. S. Ekun, Dr. P. M. Etaware and Dr. Olufunmilayo Idowu declare that they have no conflict of interest and that they actively participated in the research both in the field and in the procurement of materials for morphological and molecular analysis.

Thank you

Peter M. Etaware (Ph.D.)

## Ethical Approval

‘Not applicable’.

## Consent to Participate

All the authors gave their consent to participate in this research.

## Consent for Publication

All the authors unanimously agreed that this article should be published

## Availability of Data and Material

All data and material are present in this publication.

## Authors’ Contributions

E.V.S. and A.C.O conceptualized and designed the experiment. E.V.S conducted the research and E. P. M. wrote the draft manuscript. E. P. M. and O. O. reviewed the manuscript. All authors approved the final version of the manuscript.

